# Charting a tissue from single-cell transcriptomes

**DOI:** 10.1101/456350

**Authors:** Mor Nitzan, Nikos Karaiskos, Nir Friedman, Nikolaus Rajewsky

**Affiliations:** John A. Paulson School of Engineering and Applied Sciences, Harvard University, 29 Oxford St, Cambridge, Massachusetts 02138, USA; Broad Institute of MIT and Harvard, 415 Main St, Cambridge, Massachusetts 02142, USA; School of Computer Science and Engineering, The Hebrew University of Jerusalem, Jerusalem 9190401, Israel.; Systems Biology of Gene Regulatory Elements, Berlin Institute for Medical Systems Biology, Max Delbrück Center for Molecular Medicine in the Helmholtz Association, 10 Robert-Roessle-Str, Berlin 13125, Germany.; Institute of Life Sciences, The Hebrew University of Jerusalem, Jerusalem 9190401, Israel.

## Abstract

Massively multiplexed sequencing of RNA in individual cells is transforming basic and clinical life sciences. However, in standard experiments, tissues must first be dissociated. Thus, after sequencing, information about the spatial relationships between cells is lost although this knowledge is crucial for understanding cellular and tissue-level function. Recent attempts to overcome this fundamental challenge rely on employing additional in situ gene expression imaging data which can guide spatial mapping of sequenced cells. Here we present a conceptually different approach that allows to reconstruct spatial positions of cells in a variety of tissues without using reference imaging data. We first show for several complex biological systems that distances of single cells in expression space monotonically increase with their physical distances across tissues. We therefore seek to map cells to tissue space such that this principle is optimally preserved, while matching existing imaging data when available. We show that this optimization problem can be cast as a generalized optimal transport problem and solved efficiently. We apply our approach successfully to reconstruct the mammalian liver and intestinal epithelium as well as fly and zebrafish embryos. Our results demonstrate a simple spatial expression organization principle and that this principle (or future refined principles) can be used to infer, for individual cells, meaningful spatial position probabilities from the sequencing data alone.

## INTRODUCTION

Single cell technologies have revolutionized our understanding of the rich heterogenous cellular populations that compose tissues, the dynamics of developmental processes, and the underlying regulatory mechanisms that control cellular functions [1–4]. However, to understand the collective function and dysfunction of tissues, the intricate pathways of development, and mechanisms of cell-to-cell communication, it is crucial to have access not only to the identities of single cells but also to their spatial context and the tissue-wide expression patterns they confer. This is a challenging task since in the process of characterizing the RNA profiles of single cells by single cell RNA sequencing (scRNA-seq), the cells are commonly dissociated and their original spatial context is lost.

Considerable efforts have been undertaken to address the challenge of obtaining transcriptomic information along with spatial context [5,6]. However, current methods are still either technically challenging, expensive and not widely available, or limited in terms of spatial resolution, number of reads per cell, the number of genes whose expression can be reliably measured, the properties (e.g. length) of transcripts that can be mapped, and the types of tissues that can be analyzed.

In parallel to the aforementioned experimental efforts, two seminal papers tackled the spatial inference problem from a computational perspective [7,8]. The key idea is that cells can be identified by their expression profile over an informative set of marker genes, and therefore, a reference atlas that captures the spatial expression patterns of these marker genes can be used as a guide for assigning spatial coordinates to single cells. Provided that the reference atlas encompasses large numbers of genes, is sufficiently quantitative, and of good resolution, it is possible to combine it with a corresponding scRNA-seq dataset from the same tissue, and restore spatial expression profiles. This scheme was successfully exploited for a number of different tissues [7–13], including complete zebrafish [7] and *Drosophila* embryos [14]. However, such methodologies heavily rely on the existence of a (usually extensive) reference spatial expression database, which may not always be available, or straightforward to construct. Moreover, in practice the number of reference marker genes is rarely sufficiently large to label each spatial position with a unique combination of reference genes, making it impossible to uniquely resolve cellular positions.

Here, we present novoSpaRc (de novo Spatial Reconstruction), a novel computational framework that enables the spatial reconstruction of single-cell gene expression *de novo*, with no inherent reliance on an existing reference atlas and the flexibility to introduce prior information when it does exist (Fig. 1a). The central idea is to make simple assumptions about how gene expression is organized in space and then to find the mapping of sequenced cells to physical positions which best respect the assumption. Here, we will explore the assumption that gene expression between nearby cells is generally more similar than gene expression between cells which are separated by larger distances. We stress that this is an assumption about overall gene expression across space. Individual genes may very well have sharp expression territories from one cell to a neighboring cell. Our assumption just states that overall, expression of individual genes should only rarely look like salt and pepper patterns but should be organized, for most genes, in (gene specific) spatial territories. This assumption can be readily tested. We show that at different stages of the developmental process of organisms, or in different tissues in matured organisms, cells that are physically close are also *close* in expression space, and vice versa. Again, this occurs despite existing sharp boundaries in expression patterns for different genes, since *closeness*, which will be properly defined below, depends on the combined effect of genes composing the full transcriptome. Mathematically, we leverage this property for the reconstruction of expression patterns across tissues by aligning (a) structural similarities between the graphs generated for single cells in expression space and physical space, and potentially, if available, (b) the expression profiles of marker genes in single cells and a spatial reference atlas. We show that this can be formulated as a generalized optimal transport problem [15–17] (Fig. 1b), and specifically as an interpolation between entropically regularized Gromov-Wasserstein [18,19] and optimal transport [20] objectives. Optimal transport is a framework which has proven to be increasingly valuable for diverse fields, including biology [21,22], and specifically, makes the reconstruction task feasible even for large datasets.

We show that novoSpaRc successfully reconstructs a variety of two- and three-dimensional tissues effectively spanning one- and two-dimensional physical spaces due to intrinsic symmetries, by solely using the sequencing data and the geometric features of the physical space (which is not possible by any existing method), and it outperforms state-of-the-art methods when prior information is available. Furthermore, novoSpaRc enables identification of spatial archetypes of spatially co-expressed genes. novoSpaRc is versatile, flexible, is applicable when no prior structural or transcriptomic information is available, and is independent of the single-cell technology used. We believe that novoSpaRc offers a novel conceptual perspective on the spatial inference problem, it has the potential of being applicable to precious material where no atlases can be registered, and can be very useful in the collaborative effort to characterize various tissues in single-cell resolution (e.g. in Human Cell Atlas project [23,24]).

**Fig. 1:**
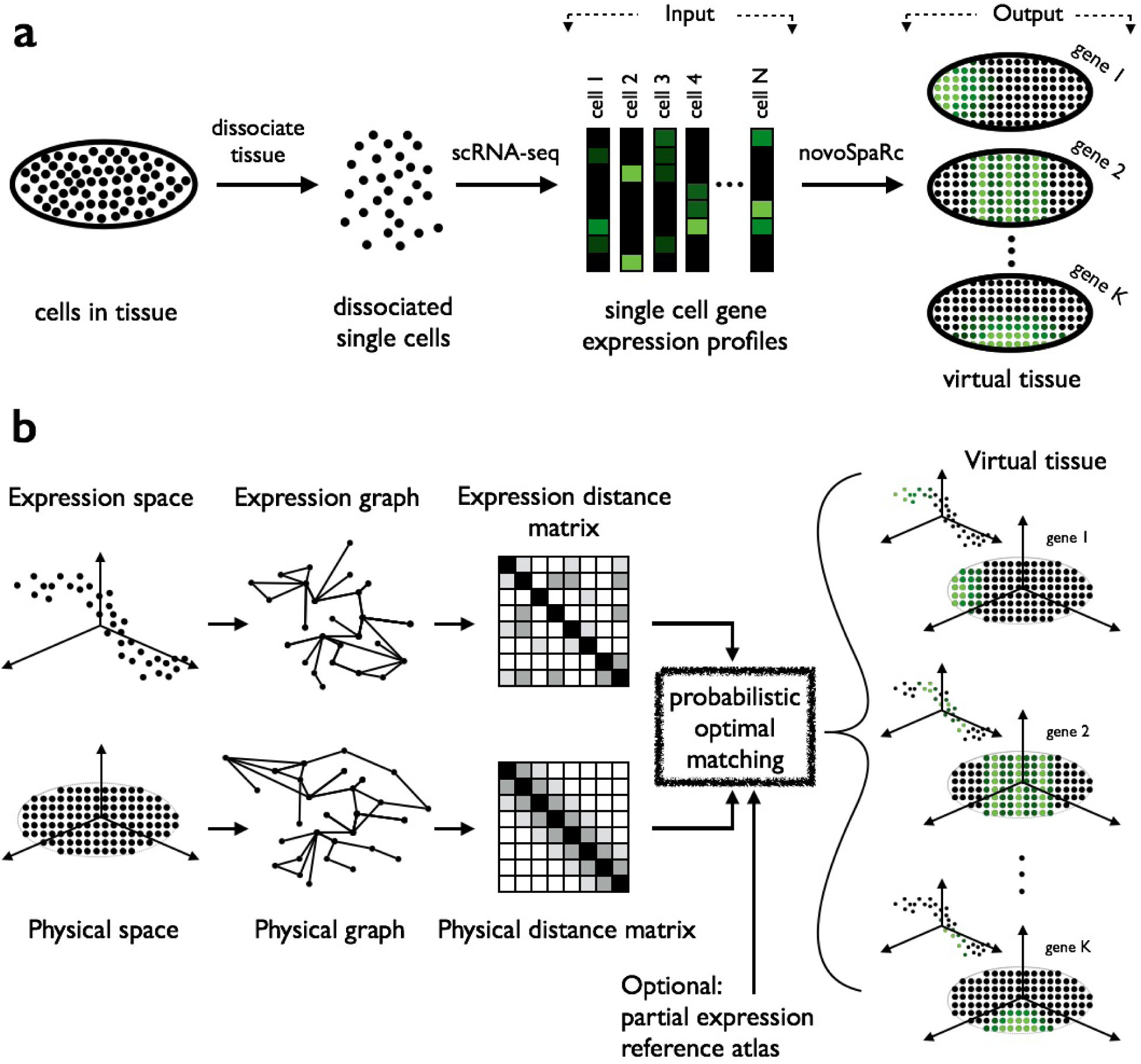
Overview of novoSpaRc.

## RESULTS

### The spatial reconstruction scheme

We start with expression profiles of single cells, all dissociated from the same tissue (usually generated by scRNA-seq). Our aim is to reconstruct the spatial patterns of expression of all genes across the tissue. Here we focus on examining stereotypical tissue patterns. Such patterns are repeated many times, either within the same tissue (e.g., crypt-to-villi axis in the small intestine) or across organisms (e.g., embryos at a specific developmental stage). Reconstructing a stereotypical pattern is important for understanding tissue structure and function. In principle our methods could be applied to the harder problem of reconstructing a specific biological instance.

We pose the problem as one of embedding single cells with their original spatial context in the canonical tissue. That is we find correspondence between two groups. The first is the *N* high-dimensional (dimensions corresponding to the number of genes *g*) expression profiles retrieved from scRNA-seq, while the second group is represented by *M* cellular locations within a tissue when such locations are known (e.g. the reproducible cellular locations in the *Drosophila* embryo during late stage 5 of development, pre-registered for ~6,000 cells [25]). For clarity, we will use *cells* when referring to expression profiles and *locations* when referring to physical cellular locations.

A canonical physical map is at times unknown. In these cases, we can flexibly convert the second group to be represented by the locations of the nodes within a regular lattice covering the shape of the original tissue (or, in fact, any distribution of finite support with predefined desirable properties). If the (effective) dimension of the tissue to be reconstructed is unknown, it can be possibly approximated by computing the intrinsic dimensionality of the manifold spanned by single cells lying in the high dimensional expression space (by using for example a maximum likelihood-based approach [26]).

The main issue is what would distinguish a biologically correct embedding from other possible ones. For a large variety of canonical tissues, cells that are *close* to each other in physical space are also *close* to each other in expression space. More generally, we hypothesize that in many cases this relationship is (at least locally) increasingly monotonic. Biologically, this phenotype can result from multiple mechanisms — gradients of morphogens and nutrients, trajectory of cell maturation, and communication between neighboring cells. While all of these can induce either smooth gradients or sharp boundaries (or combinations thereof) in gene expression patterns, as long as there are spatial shifts between sharp boundaries exhibited by different genes, our hypothesis would hold since *closeness* is a combined property of all genes in the transcriptome. We found that this is the case for the tissues we have analyzed (Fig. 2b,f and Fig. 3b).

**Fig. 2:**
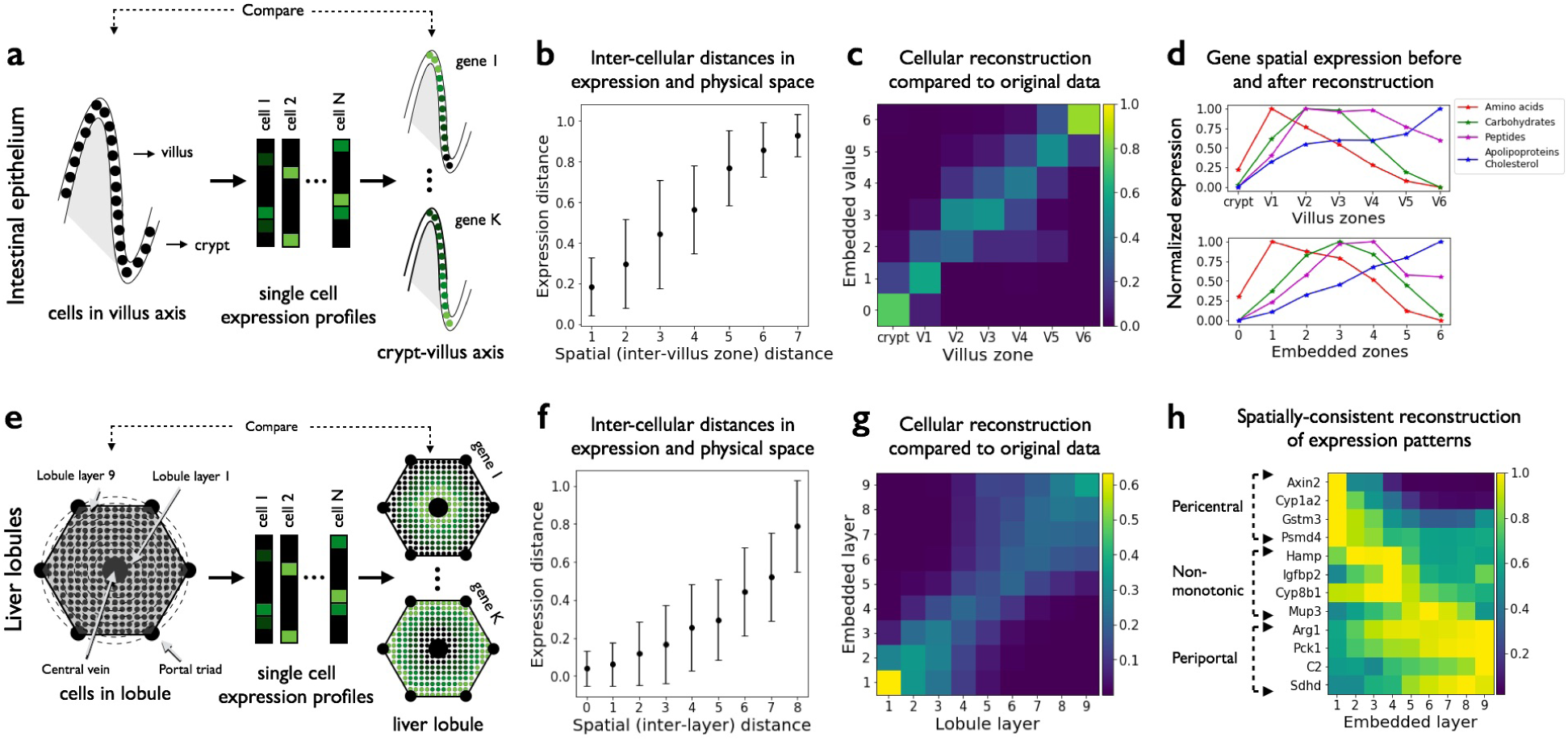
novoSpaRc reconstructs tissues with effective 1D structure. The scheme and reconstruction results are shown for the mammalian intestinal epithelium (top row, a) and the liver (bottom row, e). (b, f) There is an increasing monotonic relationship between the cellular pairwise distances in expression space and in physical space for both tissues. Error bars represent standard deviation. (c, g) novoSpaRc infers the original spatial context of single cells in both tissues with high accuracy, demonstrated by heatmaps showing the inferred distribution over embedded layers (rows) for the cells in each of the original layers (columns). In addition, the de novo embedding captures spatial division of labor of averaged expression of genes that play a role in the absorption of different nutrient classes in the intestine (d), and captures spatial expression patterns (pericentral, periportal and non-monotonic patterns) at single gene resolution in the liver (h).

**Fig. 3:**
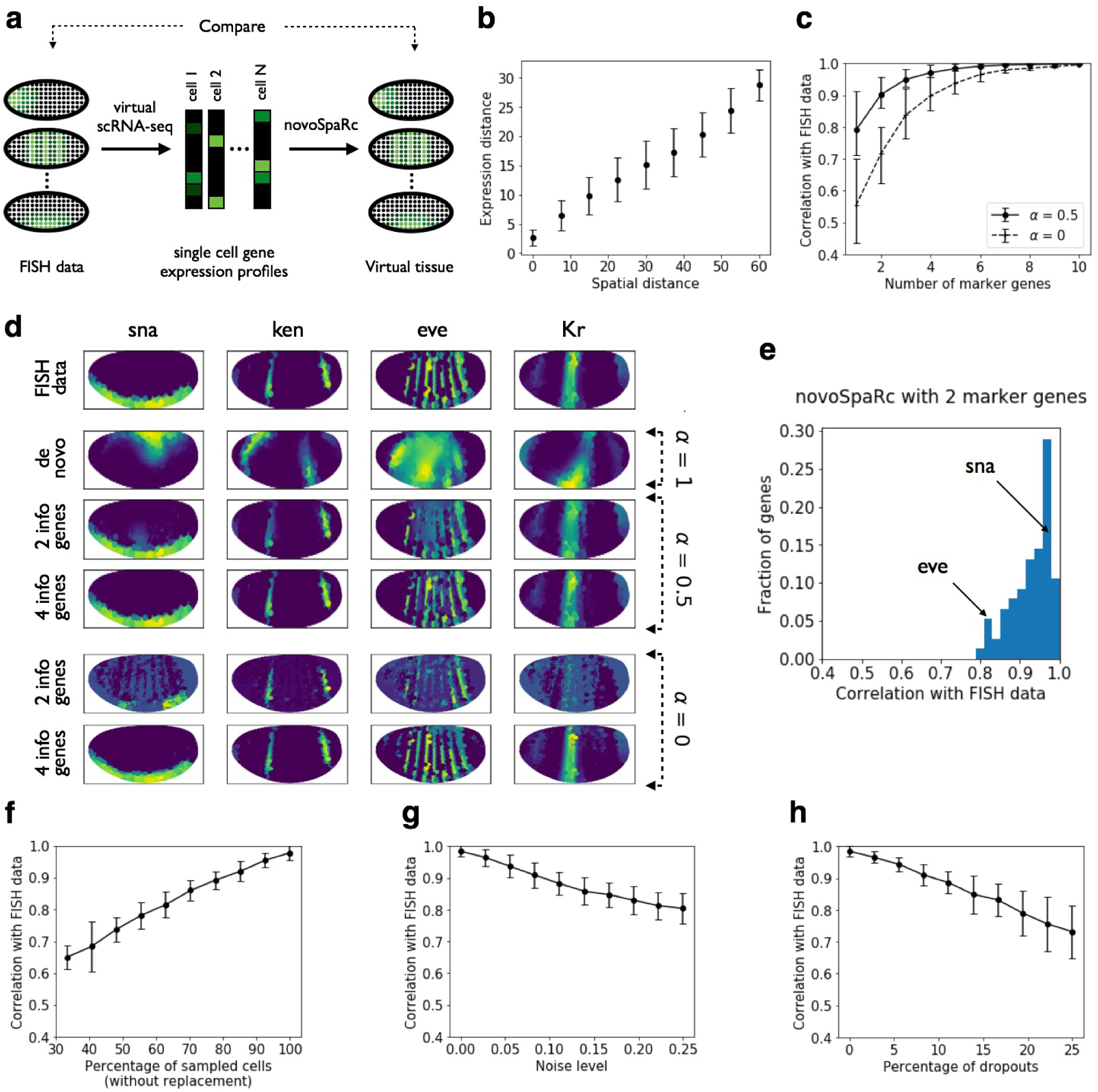
novoSpaRc reconstructs the *Drosophila* embryo based on the BDTNP dataset [25]. (a) schematically, FISH data is utilized to create virtual scRNA-seq data, which novoSpaRc then uses to reconstruct a virtual embryo. (b) There is an increasing monotonic relationship between the cellular pairwise distances in expression space and in physical space. (c) novoSpaRc reconstructs the original spatial context of single cells within the *Drosophila* embryo. The quality of reconstruction increases with the number of marker genes and saturates at perfect reconstruction at approximately 4 marker genes, when taking structural information into account (*α* = 0.5, black line). This outperforms reconstruction that relies only on marker gene information (*α* = 0, dotted line). (d) Visualization of reconstruction results for 4 genes. The original FISH data (first row) is compared to reconstruction using novoSpaRc without any marker gene information (de novo, *α* = 1), reconstruction that exploits both structural and marker gene information (*α* = 0.5), and reconstruction that uses only marker gene information (*α* = 0). When examining an instance of novoSpaRc reconstruction using 2 marker genes (*α* = 0.5), the distribution of gene-specific coefficients of correlation with the FISH data reveals that lower correlation values correspond to finer expression patterns (e). Correlation of the reconstructed expression patterns (*α* = 0.5) to the original FISH expression data increases with the percentage of sampled single cells (without replacement) (f), and steadily decreases with noise level (g) and percentage of dropouts in the data (h). Results are averaged over 100 random choices of 4 marker genes for (f-h). Error bars represent standard deviation.

With this basic assumption, we formulate an optimization problem over embedding of cells to locations. The optimization prefers embeddings that preserve pairwise distances between cells (in expression space) and the distances between their assigned locations (in physical space). This problem intuitively corresponds to finding a correspondence between the two low-dimensional manifolds in expression and physical spaces. If additional information regarding a reference expression atlas of several marker genes exists, we would like the embedding to match that information as well.

In principle, there is no reason to require a deterministic embedding of cells to locations. Instead, we pose the problem as finding a probabilistic embedding between the cells and locations. That is, each single cell will be assigned a distribution over cellular locations. A probabilistic mapping is preferable for several reasons: **(1) Single cell data does not yield an exact 1-to-1 matching problem**. (i) When a tissue is dissociated into single cells, we will generally not be able to retrieve information for the full batch of single cells, but only for a certain fraction of them, due to experimental constraints. (ii) There would generally not be information about the number of original single cells in the tissue and their exact location, meaning we would need to resort to assignment of single cells over a grid. (iii) Even in cases where there are known, reproducible cellular locations, and there is the possibility to dissociate many nearly-identical tissues to increase the single cell coverage, we would still expect to have cellular locations that correspond to multiple single cells, and cellular locations that do not correspond to any of the single cells in the dataset. **(2) Probabilistic mapping would yield smoother expression patterns and would be more robust to the noisy, partially sampled single cell data.** Given imperfect data, as is the case for experimental setups, we may be uncertain about the exact location of a dissociated single cell and would rather place it in a certain *neighborhood* of the tissue (or, probabilistically spread it over several locations in that area). This is motivated both by noise and dropouts in the original data, and the fact that if we are mapping single cells to a grid, their *true* original location may be in between several nodes (cellular locations) on the grid, in which case their *true* mapping should be distributed over the grid nodes surrounding the original location, weighted by their corresponding distance from that location. **(3) Probabilistic matching is more efficient computationally**. Intuitively, we replace a discrete optimization problem over a large combinatorial space with a continuous optimization of a smooth function, which allows us to employ more efficient optimization methods. **(4) We are interested in the reconstructed expression patterns over stereotypical tissues, and not necessarily in assigning single cells their exact original location**.

To optimize this criteria we need to set the distances to be preserved by the embedding. As the expression profiles are in high-dimensional space, metric distances are prone to multiple limitations. Instead, we use steps motivated by non-linear dimensionality reduction methods (e.g., Isomap [27]). However, at this stage we do not require finding low-dimensional coordinates, but rather constructing a robust distance matrix. For symmetry we apply the same procedure to both cells and locations independently (Fig. 1b, first column). We start by computing pairwise distances between entities. We chose as a distance metric the Euclidean distance for the physical space (locations) and the correlation-based distance for the expression space (cells), but other measures can be used. These however do not capture the true geometry of nonlinear low-dimensional manifolds. Thus, we use these pairwise distances to construct a k-nearest neighbors graph (Fig. 1b, second column). From these graphs, we compute the shortest path lengths for each pair of cells, resulting in graph-based distance matrices for cells, 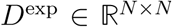, and for locations, 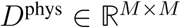 (Fig. 1b, third column).

Then, optionally, if a reference atlas is available, we compute the matrix of agreement, 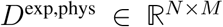, between each of the cells to each of the locations, based on the inverse correlation between the partial expression profile for each location given by the atlas and the respective expression profile for each cell.

Equipped with these measures of intra- and inter-dataset distances, we set out to find an optimal (probabilistic) assignment of each of the single cells to cellular physical locations.

We formulate this problem as an optimization problem within the generalized framework of optimal transport [15–17]. Optimal transport is a mathematical framework that was first established in the eighteenth century by Gaspard Monge and was initially motivated by a question of the optimal (minimal cost) way to rearrange one pile of dirt into a different formation (the respective minimal cost is appropriately termed earth mover’s distance). The framework evolved both theoretically and computationally [16,17,20] and drew extensions to correspondence between pairwise similarity measures via the Gromov-Wasserstein distance [18,19]. Thus, in our context, it allows us to build upon these results and tools to feasibly solve the cellular assignment problem.

We would like to find a probabilistic embedding, 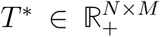, of the *N* cells to the *M* locations, which would minimize the discrepancy between the pairwise graph-based distances in expression space and in physical space, and if a reference atlas is available, minimize the discrepancy between its values across the tissue and the expression profiles of embedded single cells. For each cell *i*, the value of *T_i,j_* is the relative probability of embedding it to location *j*.

These optimization requirements over *T*\* are formulated as follows. We measure the pairwise discrepancy of *T* using the the Gromov-Wasserstein discrepancy [18]

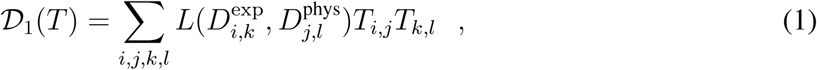

where *L* is a loss function, specifically we use the quadratic loss 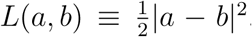. This term captures our preference to embed single cells such that their pairwise distance structure in expression space would resemble their pairwise distance structure in physical space. Intuitively, if expression profiles corresponding to cells *i* and *k* are embedded into cellular locations *j* and *l*, respectively, then the distance between *i* and *k* in expression space should *correspond* to the distance between *j* and *l* in physical space (e.g. if *i* and *k* are *close* expression-wise they should be embedded into *close* locations and vice versa). The discrepancy measure weighs these correspondences by the respective probability of the two embedding events.

To measure the match to existing prior knowledge, or an available reference atlas, we use the measure:

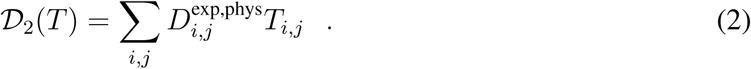

This term represents the average discrepancy between cells to locations according to the reference atlas, weighted by *T*.

Finally, we regularize *T* by preferring embeddings with higher entropy, where the entropy is defined as

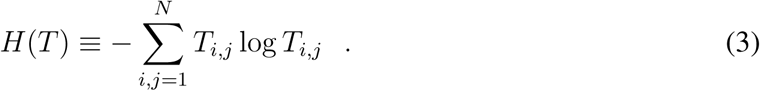

Intuitively, higher entropy implies more uncertainty in the mapping. Entropic regularization drives the solution away from arbitrary deterministic choices and was shown to be computationally efficient [20].

Putting these together, we define the optimization problem

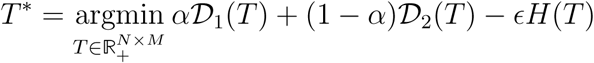

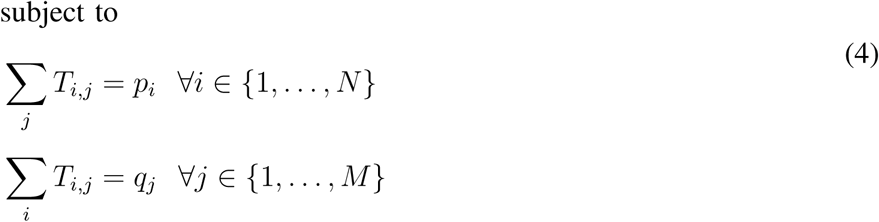

where *ϵ* is a non-negative regularization constant, and *α* ∈ [0, 1] is a constant interpolating between the first two objectives, and can be set to *α* = 1 when no reference atlas is available. The constraints reflect the fact that the transport plan should be consistent with the marginal distributions, 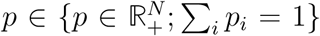 and 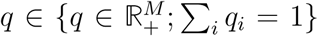, over the original input spaces of expression profiles and cellular locations, respectively. These marginals can capture, for example, varying densities of single cells in the vicinity of different cellular grid locations, or the quality of different single cell expression profiles (hence forcing low-quality single cells to have a smaller contribution to the reconstructed tissue-wide expression patterns). When such prior knowledge is lacking, *p* and *q* should be set to be uniform distributions.

We derive an efficient algorithm for this optimization problem (Methods) inspired by the combined results for entropically regularized optimal transport [20] and Gromov-Wasserstein distance-based mapping between metric-measure spaces [19].

Then, given the original single cell expression profiles, represented by a matrix 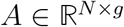, and the inferred probabilistic embedding 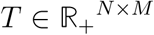, we can derive a virtual *in situ* hybridization (vISH), 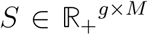, which contains the gene expression values for every cellular location of the target space:

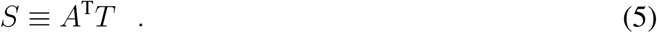

Note again that since our mapping is probabilistic, each of the cellular locations of the vISH does not correspond to a single cell in the original data. Rather, the vISH represents the expression patterns over an averaged, stereotypical tissue that the single cells could have originated from. The vISH can also be used for identifying spatially variable genes and for constructing spatial archetypes and extracting spatially related genes from scRNA-seq data, as will be described below.

#### novoSpaRc reconstructs tissues with effective 1D structure

we attempted to de novo reconstruct tissues with inherent symmetries which render them effectively 1-dimensional, with no reliance on marker gene information. We focused on scRNAseq datasets available from the mammalian intestinal epithelium [12] and the liver [9]. Schematic figures for the reconstruction process are shown in Figs. 2a and 2e, respectively.

For both tissues, cells were classified into distinct zones, or layers, based on robust marker gene information (7 zones for the intestinal tissue [12], 9 layers for the liver [9]). We found that the average pairwise distances between cells in expression space increased monotonically with the pairwise distances in physical 1-dimensional space (Fig. 2b,f), consistent with our monotonicity assumption.

We used novoSpaRc to embed the expression data into one dimension. We found that the embedded coordinates of single cells correspond, on average, to their layer or zone memberships (Fig. 2c,g), suggesting that the spatial context of single cells can be reconstructed *de novo*, without the need for a reference atlas.

Furthermore, novoSpaRc captures division of labor and spatial expression patterns within the intestine epithelium (Fig. 2d), as well as within the layers of the liver lobules (Fig. 2h), where cells in different layers of the tissue perform different tasks and exhibit different expression profiles. For example, we recover (Fig. 2d, bottom) the ordering of the expression peaks along the intestinal villi (which can be interpreted as specialized division of labor) of groups of genes that play important roles in the absorption and transportation of different nutrient groups, including apolipoproteins cholesterol, peptides, carbohydrates and amino acids (Fig. 2d, top, [12]). Finally, we correctly identify spatial expression patterns of genes in the liver exhibiting pericentral, periportal or non-monotonic profiles (Fig. 2h, [9]).

Taken together, these results demonstrate that novoSpaRc can reliably reconstruct the spatial patterns of biological tissues with effective 1-dimensional structure, with no reliance on marker gene information.

#### novoSpaRc reconstructs tissues with effective 2D structure

We next turned to spatially reconstruct a more challenging, higher dimensional object. We chose to focus on the well-studied *Drosophila* embryo. At late stage 5, the *Drosophila* embryo consists of ~6,000 cells and exhibits a bilateral symmetry. The expression levels of ~80 marker genes were registered using fluorescence in situ hybridization (FISH) for each of the 6,000 cells in a highly quantitative manner through the Berkeley Drosophila Transcription Network Project (BDTNP) project [25]. To test the performance of novoSpaRc, we first simulated scRNA-seq data, by effectively dissociating the BDTNP dataset into single cells, and then attempted to reconstruct the original expression patterns across the tissue without using information about the original FISH data (Fig. 3a). We first confirmed that the data adheres to novoSpaRc’s underlying assumptions. Similarly to the 1D datasets, we found a monotonically increasing relationship between the cell-cell pairwise distance in expression space and in physical space (Fig. 3b). We used novoSpaRc to retrieve the spatial expression patterns across the embryo, which exhibited high correlation with the original structure (Fig. 3c,d). We found that when employing novoSpaRc using both structural information and a reference atlas (*α* = 0.5), the BDTNP dataset was spatiall reconstructed faithfully, outperforming reconstruction based only on a reference atlas (*α* = 0), saturating at perfect reconstruction for approximately 4 marker genes (Fig. 3c,d). The *de novo* reconstruction (*α* = 1) correctly identified the different spatial local structures, including the dorsal-ventral and the anterior-posterior axes of the embryo, and the reconstructed configuration is highly similar to the original one (Fig. 3d, Extended Data fig. 1). In general, expression patterns that capture high-resolution details are more challenging to reconstruct (Fig. 3e, Extended Data fig. 1).

In addition to the reconstructed expression patterns (vISH), the optimal embedding also yields the embedding probability map for single cells (Extended Data Fig. 2). novoSpaRc captures the physical neighborhood single cells originated from, and as more marker genes are used for reconstruction, the embedding of single cells becomes more localized.

Generally, since de novo reconstruction is performed without any prior information that would *anchor* the cells, the reconstructed configuration is *similar* up to local transformations (reflections, rotations, translations) relative to the original configuration. However, there are features of a faithful reconstruction we can test for, as that the embedding of single cells onto the embryo is relatively localized, as we would expect for a biologically-meaningful embedding. Specifically, the mean standard deviation of embedding over cells is statistically significantly lower than that of a randomized embedding (Extended Data Fig. 3a). In addition the de novo reconstruction is robust and exhibits a linear response to perturbations. For example, the embedded expression pattern changes gradually with the entropic regularization parameter *ϵ*, both relative to the FISH data (Extended Data Fig. 4c) and relative to itself, in a self-consistent manner (Extended Data Fig. 3b).

To have an intuitive comparison against the original configuration, we used four marker genes to anchor the reconstruction. The correlation between the expression patterns across the reconstructed and the original embryo grows substantially as more genes are used to approximate the low-dimensional manifold in expression space (Extended Data Fig. 3a), and as the fraction of spatially-informative genes grows relative to spatially-noninformative genes (Extended Data Fig. 3b). The quality of the reconstructed embryo gradually increases with the number of available single cells (Fig. 3f when cells are subsampled without replacement and Extended Data Fig. 4d when sampling with replacement), and gradually decreases with increasing simulated additive expression noise (Fig. 3g) and increasing fraction of simulated dropouts in the data (Fig. 3h). Graudally deteriorated performance with noise can also be observed via an increasing optimization error (Extended Data Fig. 4e).

As an intermediate step bridging the BDTNP dataset and a raw scRNA-seq dataset, we applied novoSpaRc to spatially reconstruct the virtual embryo created *in silico* in [14], quantifying the expression of ~8,000 genes in each of the cells (Fig. 4a). In that case, while the data is derived from scRNA-seq, we have a *ground truth* embedding, and therefore, we can directly evaluate the results of our approach. novoSpaRc successfully reconstructs the virtual embryo (Fig. 4b), where as expected, the quality of reconstruction increases with the number of marker genes used for reconstruction. To account for cases with no ground truth expression patterns, we also quantified the consistency of virtual embryos reconstructed by using different sets of marker genes, by calculating the average pairwise Pearson correlation within such reconstructed multi-dimensional expression patterns. This consistency score indeed points to successful reconstruction and, again, increases with the number of marker genes (Extended Data Fig. 3c).

**Fig. 4:**
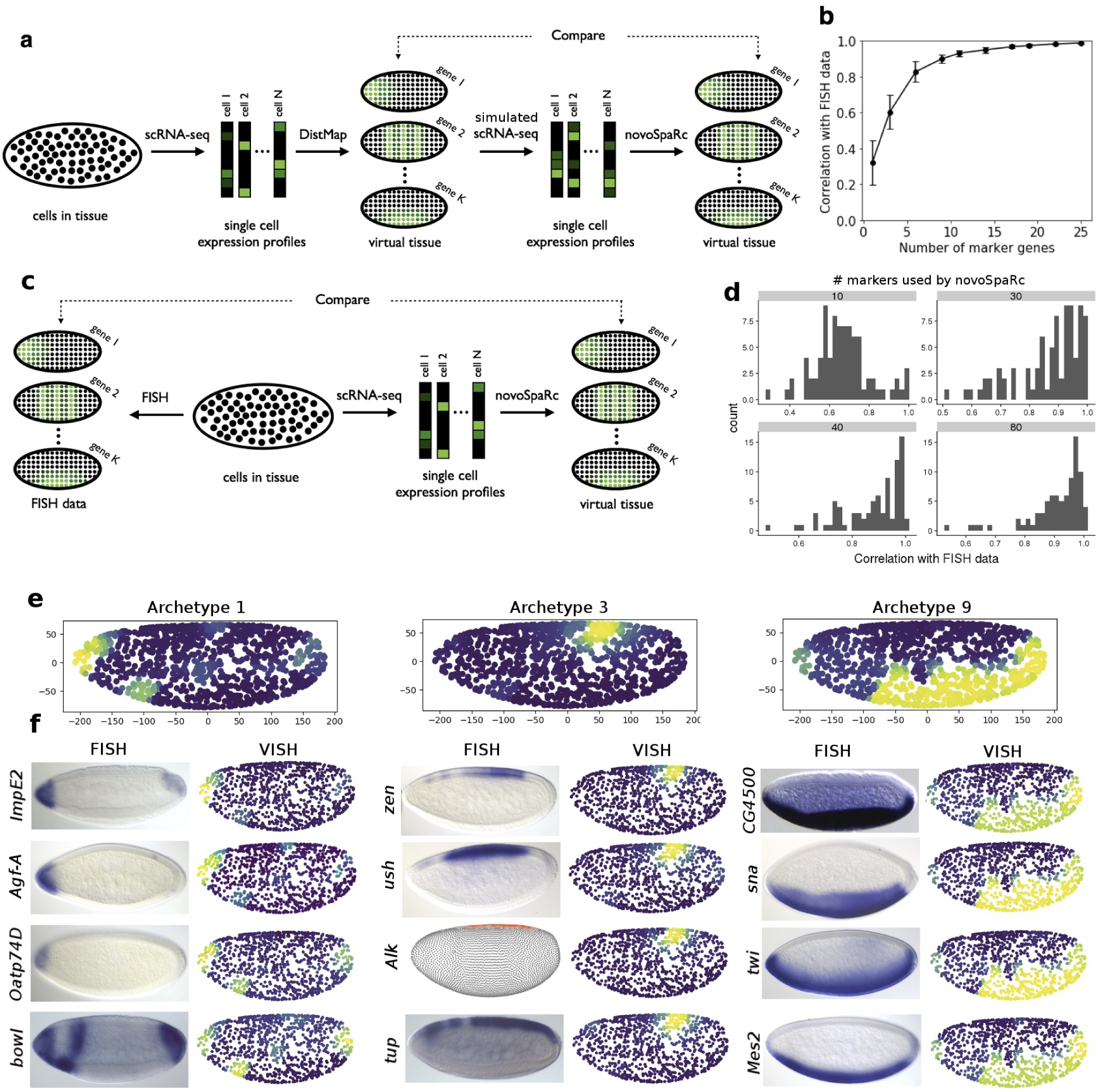
novoSpaRc reconstructs the *Drosophila* embryo based on single cell RNA-seq data. (a) Overview of the process of spatial reconstruction of the *Drosophila* virtual embryo. (b) The correlation of the reconstructed expression patterns of the virtual embryo with FISH data increases with the number of marker genes used for reconstruction. (c) Overview of spatial reconstruction of the *Drosophila* embryo using *raw* single-cell data. (d) Histograms of correlation coefficients of reconstructed expression patterns to FISH data for different genes show how the accuracy of novoSpaRc increases with the number of marker genes used. (e) novoSpaRc identifies distinct spatial domains as archetypes in the *Drosophila* embryo. Preferred spatial positioning is denoted by coloring ranging from blue (low) to yellow (high). (f) Comparison of novoSpaRc predictions (vISH) against FISH images, taken from the BDGP database [28]. For *Alk* an image was not available and DVEX was used instead [14]. Four representative genes are depicted for each spatial archetype.

The above results show that novoSpaRc is able to reconstruct the spatial information of 2-dimensional tissues with very few marker genes needed and recognizes spatial structures even without using prior information about any such marker genes.

We next attempted to reconstruct the *Drosophila* embryo of stage 6 by applying novoSpaRc to the scRNA-seq dataset generated in [14] (Fig. 4c). In that work, 84 informational genes were required for reconstruction, as well as a new computational algorithm method that distributed the ~1,300 cells over the 3,000 embryonic locations. Since novoSpaRc naturally exhibits a probabilistic mapping, we reasoned that the above dataset is a good candidate for testing its efficacy. When using both structural information and a reference atlas, we found again that the accuracy of reconstruction by novoSpaRc increases with the number of marker genes used for the atlas (Fig. 4d). *de novo*, atlas-free reconstruction by novoSpaRc of the stage 6 fly embryo worked surprisingly well. By querying the resulting vISH, we verified that novoSpaRc accurately separated the major spatial domains (mesoderm, neurogenic ectoderm, dorsal ectoderm) after gastrulation, as well as finer spatial domains (Fig. 4c-f, Extended Data Fig. 5, Methods). By clustering the highly variable genes at the vISH level into archetypes, novoSpaRc identifies spatial domains, such as the dorsal ectoderm and the mesoderm (Fig. 4e,f, Methods). The spatial archetypes can be queried for representative genes. We compared the vISHs constructed via novoSpaRc with FISH images to visually assess the accuracy of the spatial reconstruction (Fig. 4f, Extended Data Fig. 5). The patterns of genes expressed through the anterior-posterior or the dorsal-ventral axis were largely recapitulated: typical dorsal ectoderm genes such as *zen* and *ush* were co-localized dorsally (Fig. 4e,f, middle) and typical mesoderm genes such as *twi* and *sna* were co-localized ventrally (Fig. 4e,f, left). Less extensive spatial domains were reconstructed with diverse degrees of accuracy (Extended Data Fig. 5). Archetype 5, for instance, is a subdomain of the mesoderm characterized by the transcription factor *gcm*. novoSpaRc accurately grouped genes expressed in this domain, but localized the cluster in the posterior axis, although still within the mesodermic region (Extended Data Fig. 5). While capturing the main structures of the fly embryo, novoSpaRc did not capture the fine details of the expression patterns of ubiquitous genes or pair-rule genes (*eve*, *prd*) (Extended Data Fig. 5, Archetype 8).

As a second application of novoSpaRc to 2-dimensional tissues, we reconstructed the zebrafish embryo dataset of Satija et. al. [7] (Fig. 5a). Aiming for similar resolution as in [7], we mapped the single-cell transcriptomes onto a semi-circle of 64 distinct locations. The novoSpaRc reconstruction captured a good portion of the spatial signal with only a few marker genes and the overall accuracy of the reconstruction increased with the number of marker genes (Fig. 5b,c), exhibiting high correspondence between the predicted and the experimentally verified expression patterns (Fig. 5a). Moreover, novoSpaRc required fewer marker genes for comparable reconstruction results (Fig. 5a,b), no data imputation as in [7] or other specialized preprocessing, and the computational time was substantially shorter.

**Fig. 5:**
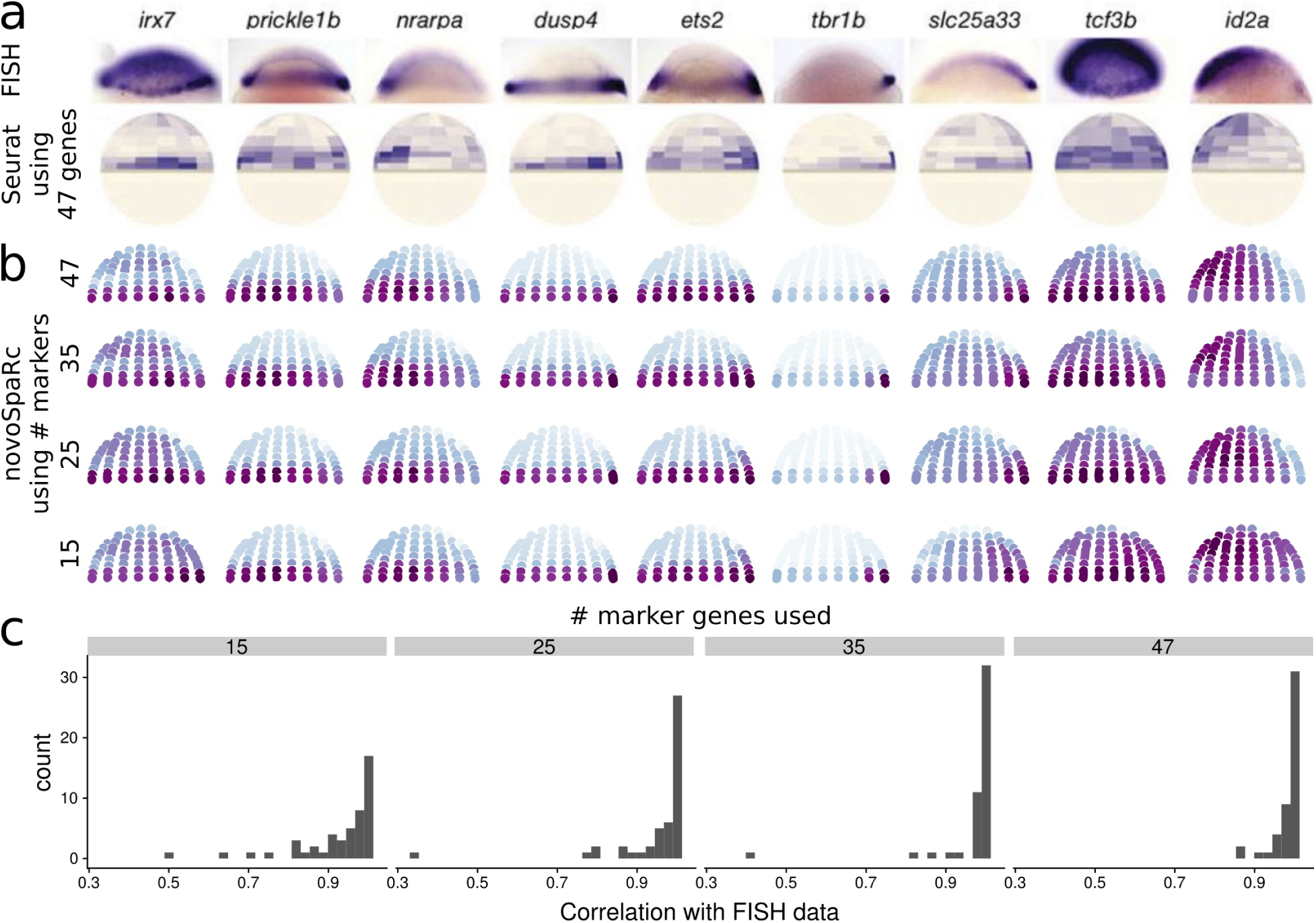
novoSpaRc reconstructs the zebrafish embryo. (a) Spatial expression patterns of 9 genes as shown by FISH experiments and as predicted by Seurat by using 48 marker genes (adapted from [7]). (b) *De novo* predictions (*α* = 1) of the expression patterns of corresponding genes by novoSpaRc for three different target space resolutions, and novoSpaRc predictions (*α* = 0.5) with 18 marker genes. (c) Histograms assessing the increase in the accuracy of novoSpaRc reconstruction with increasing number of marker genes.

Taken together, the above demonstrates that novoSpaRc is able to extract spatial information from scRNA-seq datasets and can efficiently reconstruct 2-dimensional tissues or embryonic organisms.

## DISCUSSION

In this paper, we show that relational structural information, connecting single cell expression profiles in expression space and embedded physical locations can be utilized to reconstruct expression patterns across tissues or whole organisms given single cell data, with no additional marker gene information or a spatial reference atlas. We show that this can be achieved via novoSpaRc, an efficient computational framework relying on the theory of optimal transport, that allows to smoothly interpolate between using as input only relational structural information, only a reference atlas, or both. We showed that novoSpaRc successfully reconstructs spatial expression patterns and identifies spatial archetypes across different cell types, tissues and whole organisms that are diverse in terms of their structure, function and dimensionality.

If the precise cellular locations the single cells originated from in the tissue are unknown, then the embedding can be performed onto a grid whose resolution is determined by the user. For the intestinal epithelium data, varying the grid resolution to include either less or more embedded zones than the original laser-capture-based villi zones [12] did not seem to compromise the quality of the reconstructed expression patterns (Extended Data Fig. 6). However, this generally may not always be the case. We provide several measures to reason about preferred embedded resolution and in general, to evaluate the output reconstruction results.

novoSpaRc is computationally suitable for datasets composed of a large number of single cells, and on the other hand, is robust enough to handle small datasets. In addition, single cell embedding can be done to any physical target shape and it efficiently tackles one-to-many (or many-to-one) cases, where the number of single cell expression profiles is smaller than the number of embedded cellular locations, or vice versa. Finally, the framework of novoSpaRc can flexibly accommodate prior knowledge regarding the target shape or known expression patterns of marker genes, which tend to improve the results substantially. On the other hand, when no such prior knowledge is available, although the archetypal expression patterns across tissues can be reconstructed, fine details may be difficult to discern. Future work is needed to adjust this framework to complex, non-canonical tissues with locally non-smooth expression patterns.

We expect novoSpaRc to be valuable in the study of developmental pathways and resulting expression patterns, the characterization of cellular microenvironments and their change under different conditions, and to probe cell-cell interactions.

## METHODS

### Data acquisition and analysis

The single cell RNA-seq datasets were acquired from the GEO database with the following GEO accession numbers: GSE99457 for the intestinal epithelium [12], GSE95025 for the *Drosophila* embryo [14]. The BDTNP dataset was downloaded directly from the BDTNP web-page [25].

For the cases where normalized data was not available, we adopted the standard library size normalization in log-space, e.g. if *d_ij_* represents the raw count for gene *i* in cell *j*, we normalized it as 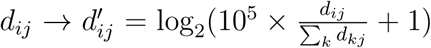. Highly variable genes were identified by plotting the dispersion of a gene as a function of its mean and selecting the outliers above cutoff values (usually 0.125 for the mean and 1.5 for the dispersion).

### Single cell embedding using optimal transport

As was discussed in the main text and Eq. 4, the optimal probabilistic coupling 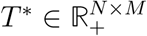 between *N* single cell expression profiles and *M* cellular locations can be framed as the solution to the following optimization problem:

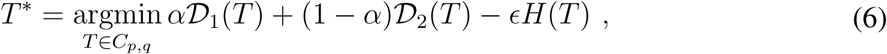

where

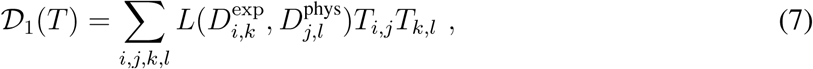

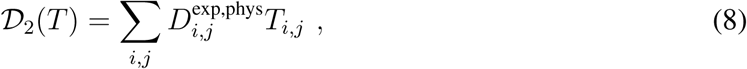

and the set of coupling between the distribution over expression profiles, 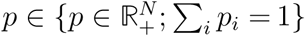, and the distribution over locations, 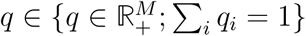, is

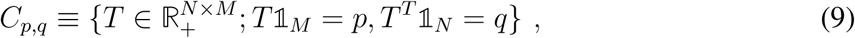

where 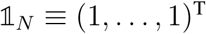.

To retrieve the coupling *T*\*, we extend upon the results for entropically regularized optimal transport [20] and Gromov-Wasserstein distance-based mapping between metric-measure spaces [19], and use projected gradient descent, where the projection is based on the Kullback-Leibler (KL) metric. Each iteration of the projected exponentiated gradient method consists of two steps; in the first step the current estimate of *T* is updated by exponentiated gradient descent step, similarly to [19], to yield 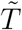:

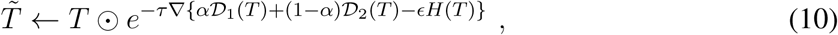

where ⊙ is an element-wise multiplication, *e*^(*x*)^ is element-wise operation, and *τ* > 0 is a small step size.

In the second step, 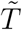 is projected back into the set *C_p_*_,*q*_ according to the KL metric:

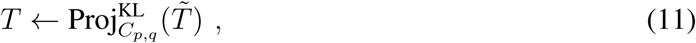

where the KL projection is

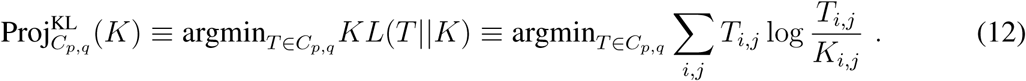

It was shown in [29] that the KL projection can be rewritten as an instance of entropically-regularized optimal transport:

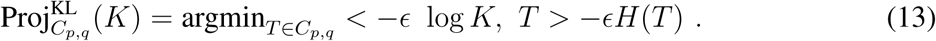

The gradient of the objective function can be written as

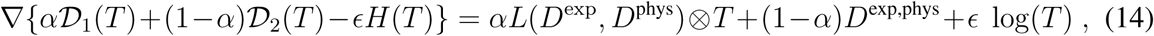

where log(*x*) is an element-wise operation, and the tensor product is defined as 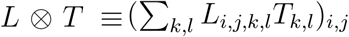.

Combining eqs. 13 and 14,

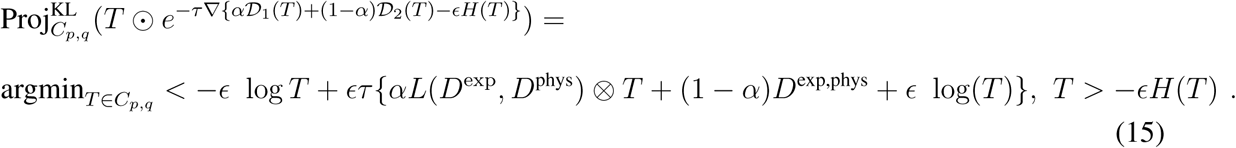

Therefore, if we set *τ* = 1/*ϵ*, each iteration of the algorithm can be simplified to a Sinkhorn projection,

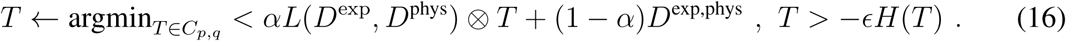

Each of the iteration steps in Eq. 16 can be computed using Sinkhorn’s fixed point algorithm [20]. Specifically,

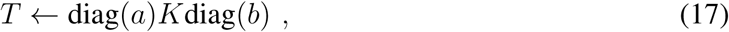

where the Gibbs kernel associated with {*αL*(*D*^exp^, *D*^phys^) ⊗ *T* + (1 − *α*)*D*^exp,phys^} is 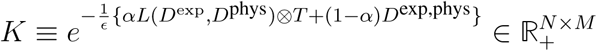. Finally, 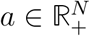 and 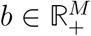 can be computed using Sinkhorn’s fixed point iterations [30] involving element-wise division:

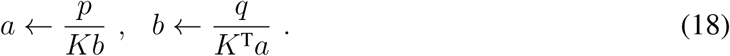

To implement the algorithm described above we based our code on a modification of the Python Optimal Transport package (https://pot.readthedocs.io/).

### Identification of spatial archetypes

The identification of spatial archetypes is performed by clustering the spatial expression of a given set of genes. The gene expression is first clustered by hierarchical clustering at the vISH level, although in principle different clustering methods can be used. The number of archetypes is then set by visually inspecting the resulting dendrogram. The expression values of each gene of the cluster are then averaged per location to produce the spatial archetype for that cluster. Representative genes for each cluster are identified by computing the Pearson correlation of each gene within the cluster against the spatial archetype. The derivation of the spatial archetypes depends strongly on the set of genes used. We observed that the set of highly variable genes generally resulted in sensible spatial archetypes.

## CODE AVAILABILITY

Code will be made available upon request.

## ACKNOWLEDGEMENTS

We thank all members of our labs for valuable comments. This work was supported by the Israeli Science Foundation, through the I-CORE program (NF) and an Alexander von Humboldt Foundation Research Award (NF). MN is supported by the James S. McDonnell Foundation, Eric and Wendy Schmidt Fund for Strategic Innovation, Israel Council for Higher Education, and the John Harvard Distinguished Science Fellows Program within the FAS Division of Science of Harvard University.

---

**Extended Data Fig. 1:**
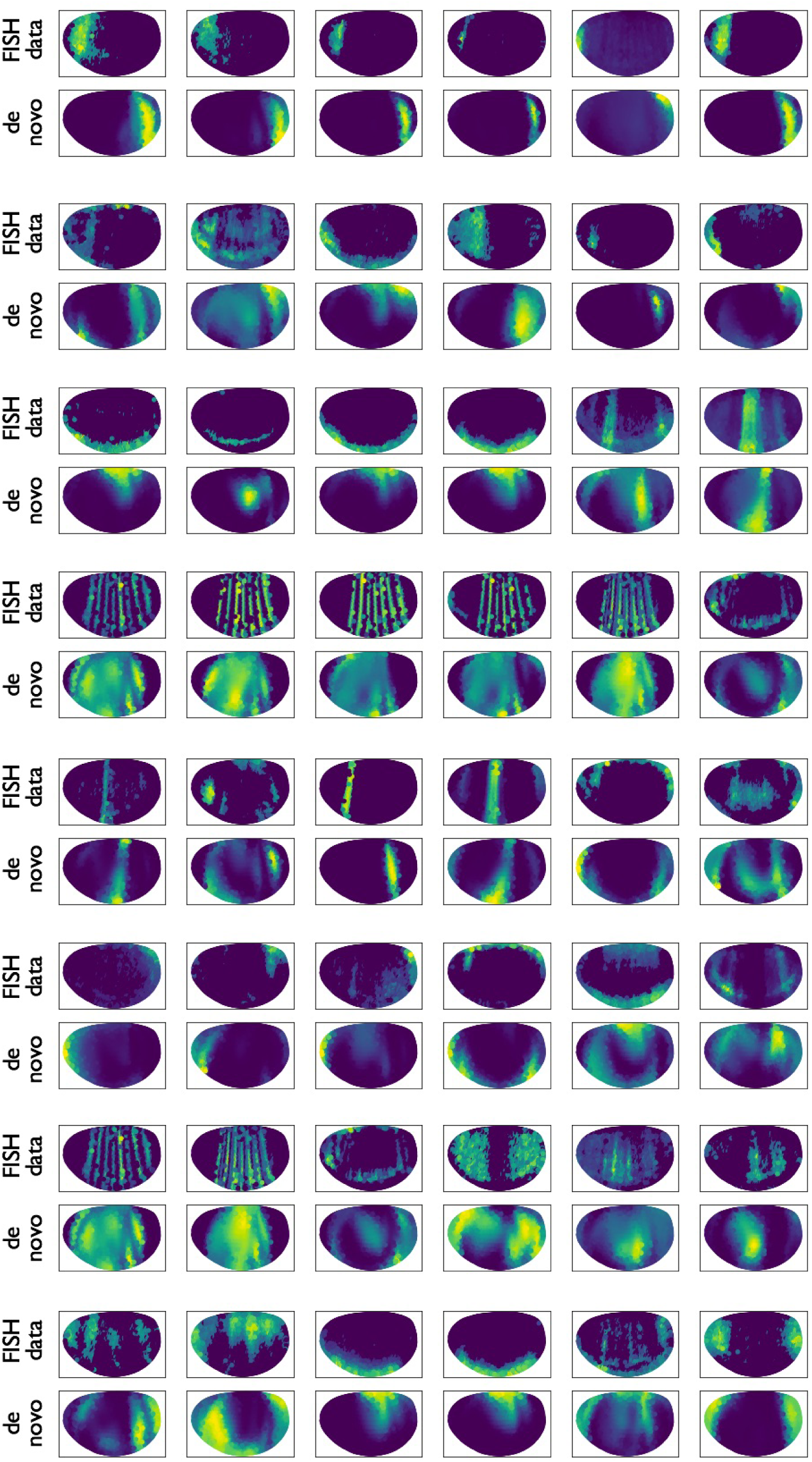
novoSpaRc reconstructs the *Drosophila* embryo de novo based on the BDTNP dataset [25]. Examples of marker gene expression patterns across the embryo comparing the original (FISH) data and the reconstructed (de novo vISH) data.

**Extended Data Fig. 2:**
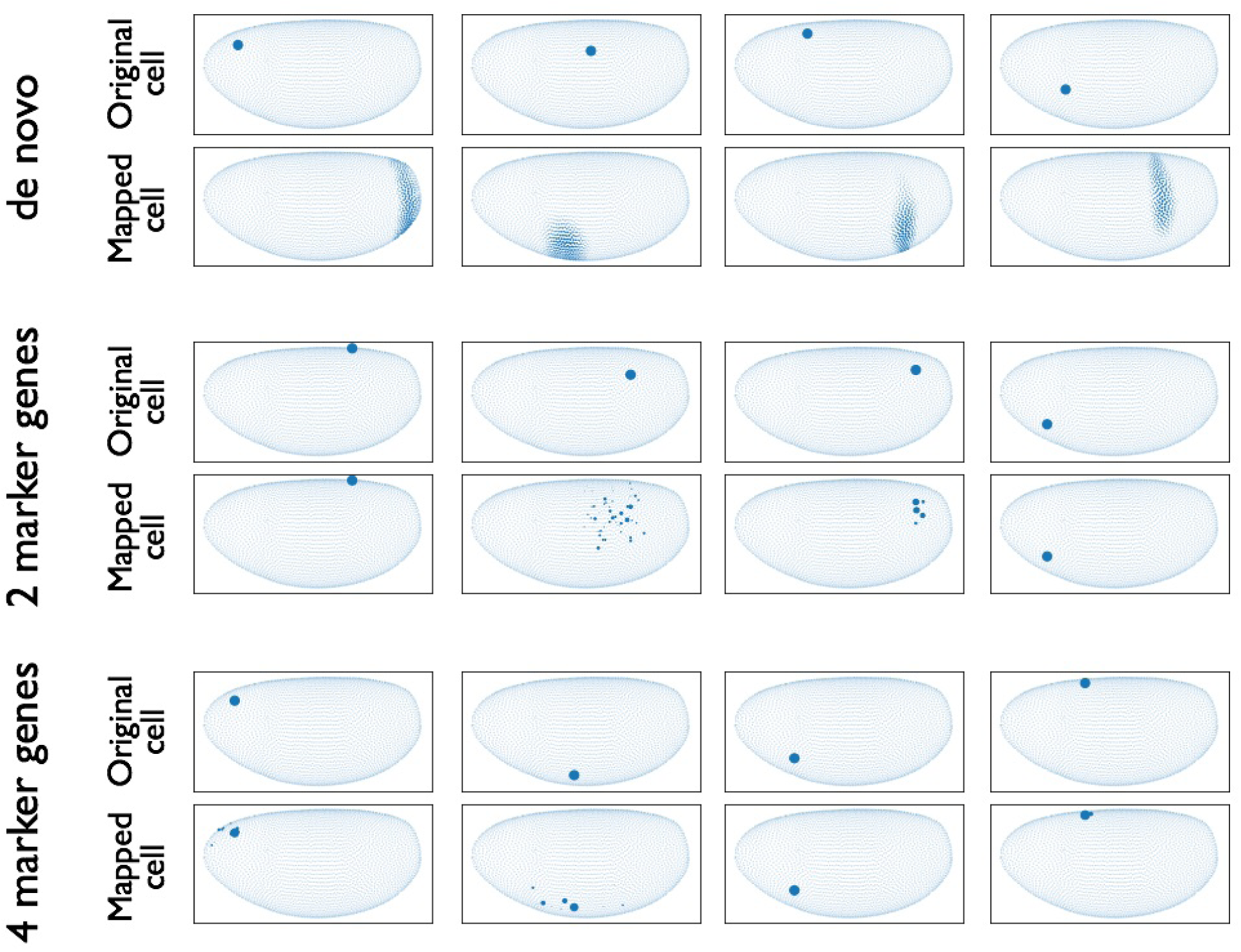
Single cell embedding produced by novoSpaRc for the *Drosophila* embryo based on the BDTNP dataset [25]. Examples of induced embeddings of single cells when reconstruction is done de novo (*α* = 1, top panel), based on two marker genes (*α* = 0.5, middle panel), or based on four marker genes (*α* = 0.5, bottom panel).

**Extended Data Fig. 3:**
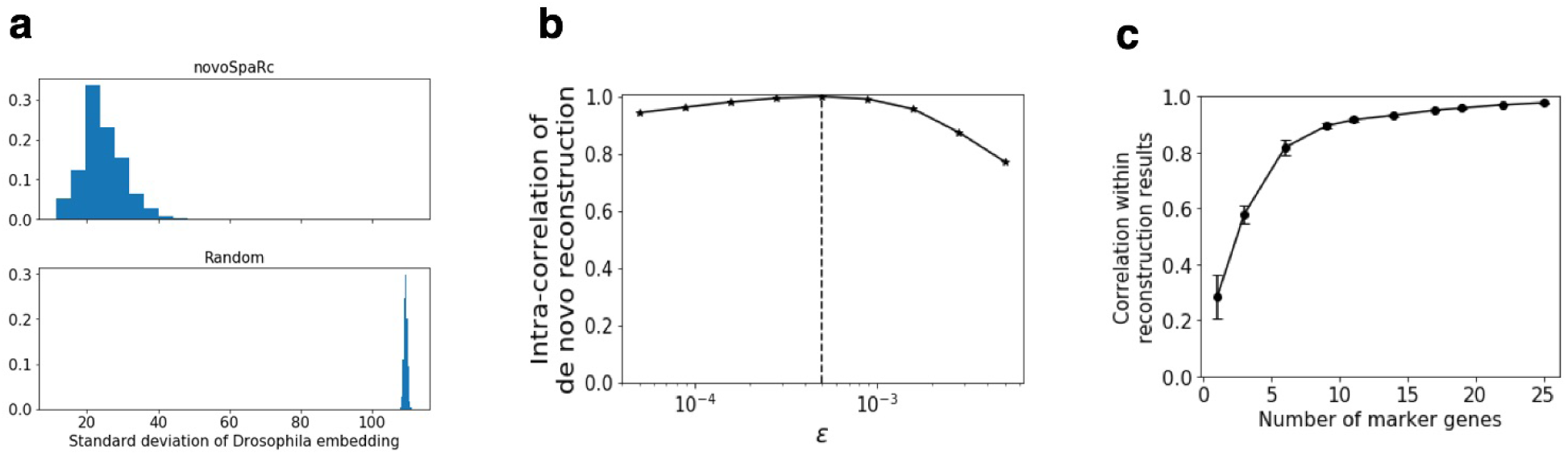
Self-consistency analysis of reconstruction with novoSpaRc. (a) The spatial standard deviation of embedded cells of the BDTNP dataset via de novo novoSpaRc (*α* = 1) vs. randomized embedding. (b) Correlation of embedded de novo expression patterns of the BDTNP dataset for different *ϵ* values with the expression pattern for *ϵ* = 5 * 10^−5^ (vertical dotted line). (c) The self-consistency of reconstruction (*α* = 0.5) of the virtual embryo increases with the number of marker genes. The consistency score was calculated as the average pairwise Pearson correlation within reconstructed expression patterns for different sets of marker genes.

**Extended Data Fig. 4:**
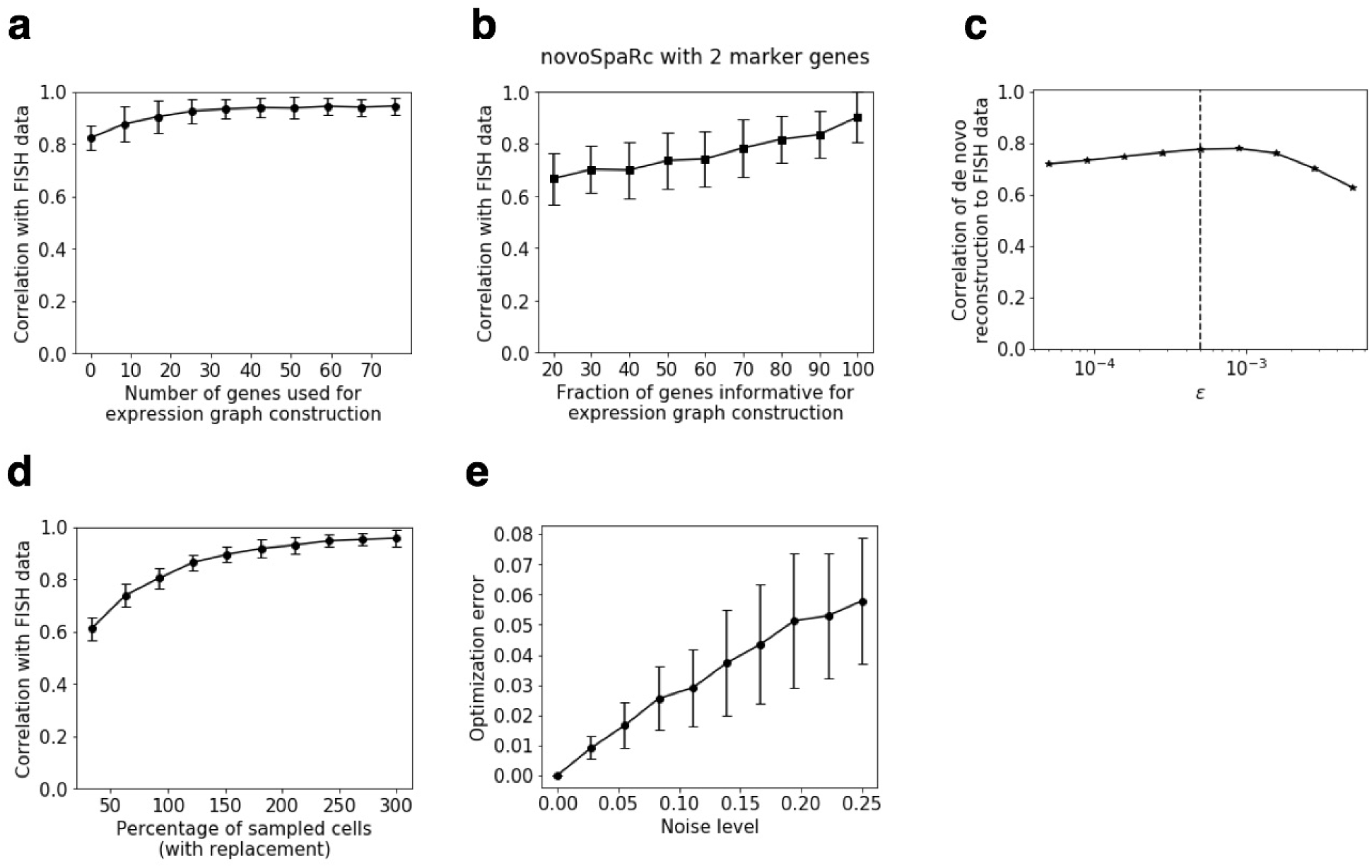
novoSpaRc reconstruction of the *Drosophila* embryo based on the BDTNP dataset is robust. Correlation of the reconstructed expression patterns (*α* = 0.5) to the original FISH expression data (a) increases with the number of genes used to construct the cellular graph in expression space, and (b) with the fraction of spatially-informative genes. (c) Correlation of the de novo reconstructed expression patterns (*α* = 1) to the original FISH data varies gradually with the entropic regularization parameter *ϵ*. (d) Correlation of the reconstructed expression patterns (*α* = 0.5) to the original FISH expression data increases with the percentage of sampled single cells (with replacement). (e) The mean value and variance of the optimization objective function (which we aim to minimize) increases with noise level. Reconstruction in (a, c-e) is performed with information for four marker genes. Reconstruction in (b) is performed with information for two marker genes. Results are averaged (and standard deviation is shown) over 100 choices of the marker genes.

**Extended Data Fig. 5:**
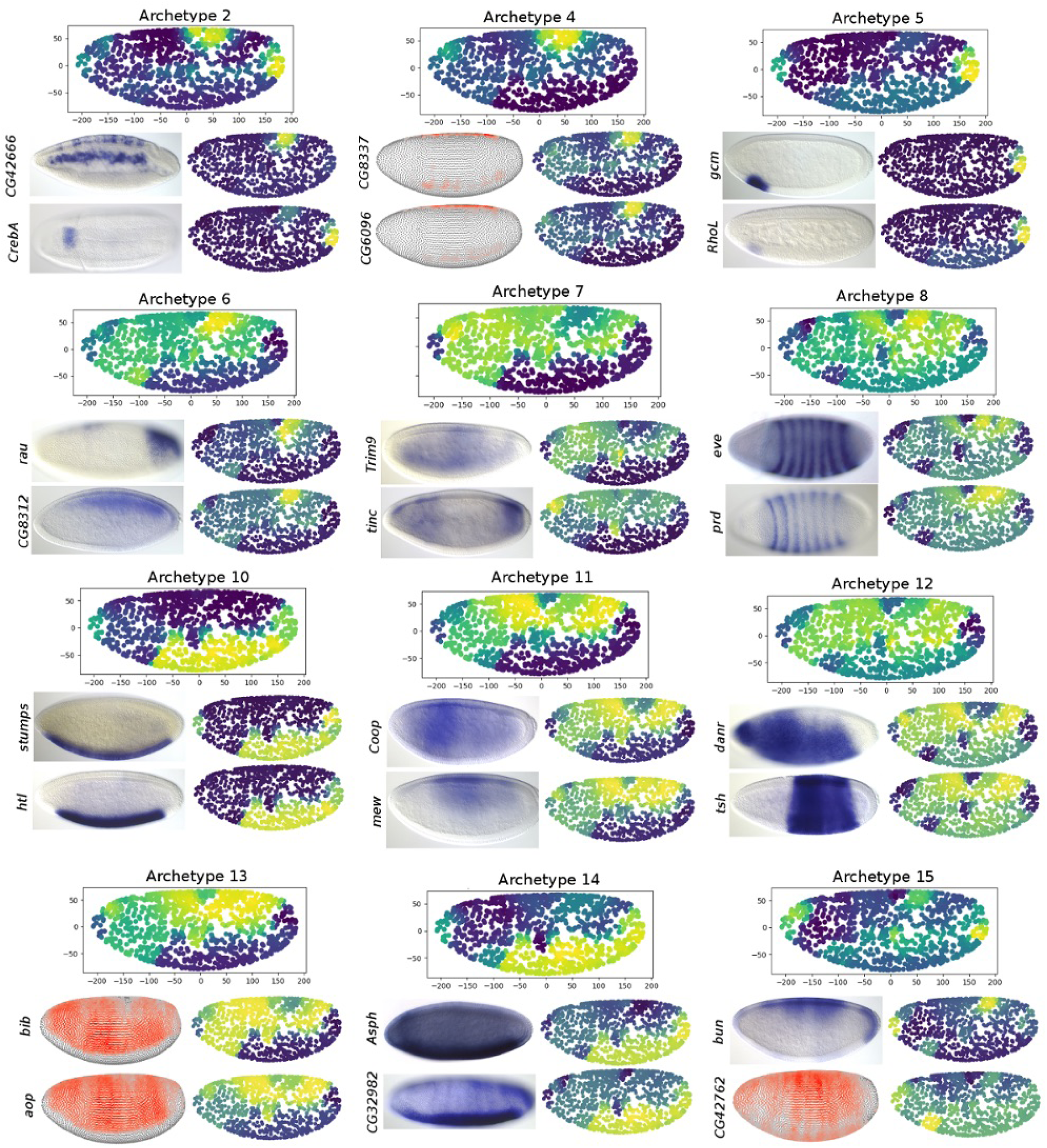
novoSpaRc identifies spatially informative archetypes using single cell RNA-seq data for the *Drosophila* embryo. The archetypes shown complement those of Fig. 4 in the main text. Preferred spatial positioning is denoted by coloring ranging from blue (low) to yellow (high). FISH images were taken from the BDGP database [28]. For genes for which an image was not available, DVEX was used instead [14]. Two representative genes are shown for each spatial archetype.

**Extended Data Fig. 6:**
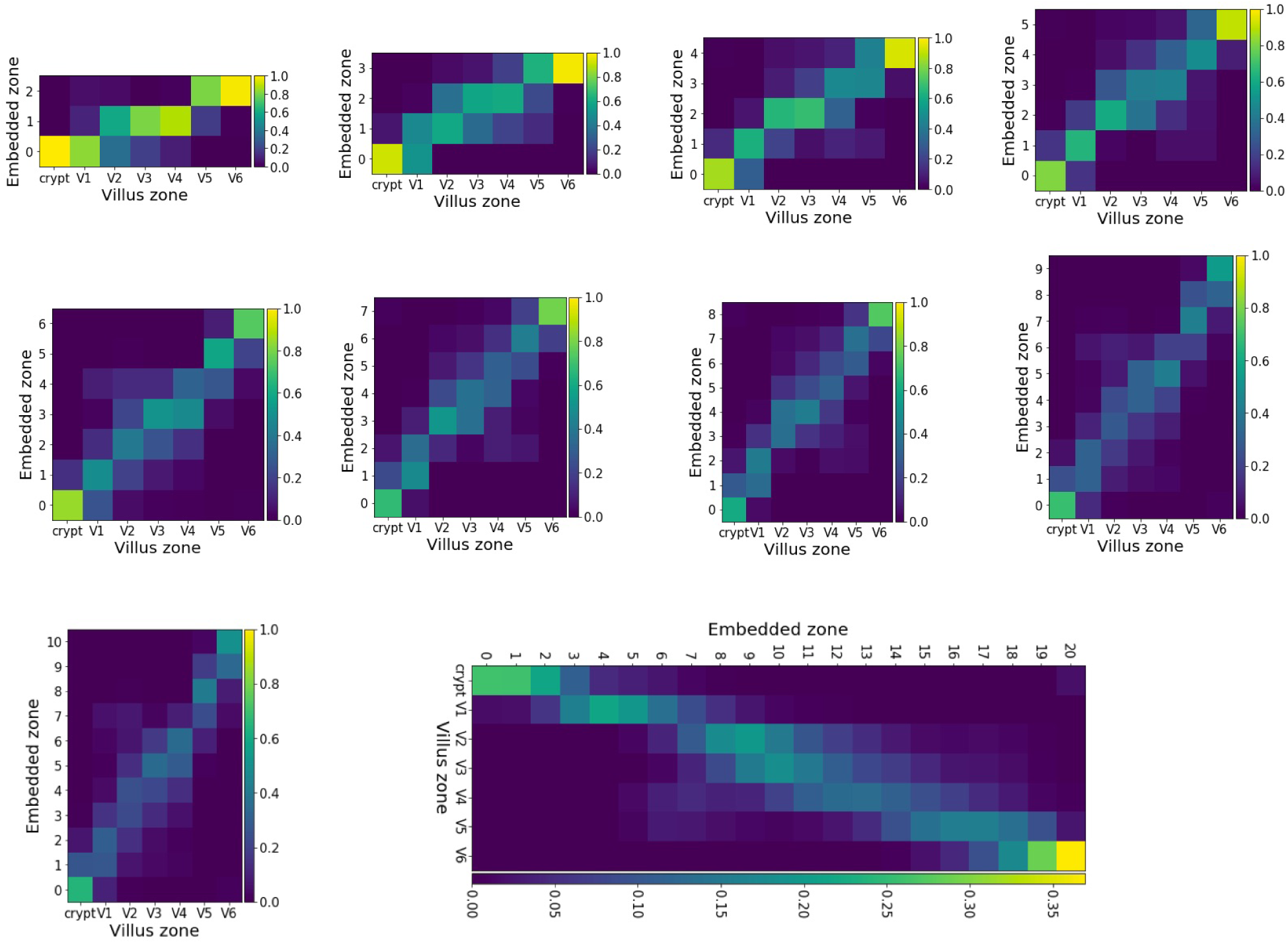
novoSpaRc reconstruction of the intestinal epithelium dataset [12] is robust and consistent for different numbers of embedded zones.

## REFERENCES

[1] E. Shapiro, T. Biezuner, and S. Linnarsson, “Single-cell sequencing-based technologies will revolutionize whole-organism science,” Nature Reviews Genetics, vol. 14, no. 9, p. 618, 2013.

[2] A. Wagner, A. Regev, and N. Yosef, “Revealing the vectors of cellular identity with single-cell genomics,” Nature biotechnology, vol. 34, no. 11, p. 1145, 2016.

[3] S. J. Altschuler and L. F. Wu, “Cellular heterogeneity: do differences make a difference?” Cell, vol. 141, no. 4, pp. 559–563, 2010.

[4] A. A. Kolodziejczyk, J. K. Kim, V. Svensson, J. C. Marioni, and S. A. Teichmann, “The technology and biology of single-cell rna sequencing,” Molecular cell, vol. 58, no. 4, pp. 610–620, 2015.

[5] N. Crosetto, M. Bienko, and A. Van Oudenaarden, “Spatially resolved transcriptomics and beyond,” Nature Reviews Genetics, vol. 16, no. 1, p. 57, 2015.

[6] E. Lein, L. E. Borm, and S. Linnarsson, “The promise of spatial transcriptomics for neuroscience in the era of molecular cell typing,” Science, vol. 358, no. 6359, pp. 64–69, 2017. [Online]. Available: http://science.sciencemag.org/content/358/6359/64

[7] R. Satija, J. A. Farrell, D. Gennert, A. F. Schier, and A. Regev, “Spatial reconstruction of single-cell gene expression data,” Nature biotechnology, vol. 33, no. 5, p. 495, 2015.

[8] K. Achim, J.-B. Pettit, L. R. Saraiva, D. Gavriouchkina, T. Larsson, D. Arendt, and J. C. Marioni, “High-throughput spatial mapping of single-cell rna-seq data to tissue of origin,” Nature biotechnology, vol. 33, no. 5, p. 503, 2015.

[9] K. B. Halpern, R. Shenhav, O. Matcovitch-Natan, B. Tóth, D. Lemze, M. Golan, E. E. Massasa, S. Baydatch, S. Landen, A. E. Moor et al., “Single-cell spatial reconstruction reveals global division of labour in the mammalian liver,” Nature, vol. 542, no. 7641, p. 352, 2017.

[10] R. Durruthy-Durruthy, A. Gottlieb, B. H. Hartman, J. Waldhaus, R. D. Laske, R. Altman, and S. Heller, “Reconstruction of the mouse otocyst and early neuroblast lineage at single-cell resolution,” Cell, vol. 157, no. 4, pp. 964–978, 2014.

[11] J. Waldhaus, R. Durruthy-Durruthy, and S. Heller, “Quantitative high-resolution cellular map of the organ of corti,” Cell reports, vol. 11, no. 9, pp. 1385–1399, 2015.

[12] A. E. Moor, Y. Harnik, S. Ben-Moshe, E. E. Massasa, K. B. Halpern, and S. Itzkovitz, “Spatial reconstruction of single enterocytes uncovers broad zonation along the intestinal villus axis,” bioRxiv, p. 261529, 2018.

[13] N. Habib, Y. Li, M. Heidenreich, L. Swiech, I. Avraham-Davidi, J. J. Trombetta, C. Hession, F. Zhang, and A. Regev, “Div-seq: Single-nucleus rna-seq reveals dynamics of rare adult newborn neurons,” Science, vol. 353, no. 6302, pp. 925–928, 2016.

[14] N. Karaiskos, P. Wahle, J. Alles, A. Boltengagen, S. Ayoub, C. Kipar, C. Kocks, N. Rajewsky, and R. P. Zinzen, “The drosophila embryo at single-cell transcriptome resolution,” Science, vol. 358, no. 6360, pp. 194–199, 2017. [Online]. Available: http://science.sciencemag.org/content/358/6360/194

[15] G. Monge, “Mémoire sur la théorie des déblais et des remblais,” Mém. de l’Ac. R. des Sc., pp. 666–704, 1781.

[16] C. Villani, Optimal transport, ser. Grundlehren der Mathematischen Wissenschaften [Fundamental Principles of Mathematical Sciences]. Springer-Verlag, Berlin, 2009, vol. 338.

[17] C. Villani, Topics in optimal transportation. American Mathematical Soc., 2003, no. 58.

[18] F. Mémoli, “On the use of gromov-hausdorff distances for shape comparison,” 2007.

[19] G. Peyré, M. Cuturi, and J. Solomon, “Gromov-wasserstein averaging of kernel and distance matrices,” in International Conference on Machine Learning, 2016, pp. 2664–2672.

[20] M. Cuturi, “Sinkhorn distances: Lightspeed computation of optimal transport,” in Advances in Neural Information Processing Systems, 2013, pp. 2292–2300.

[21] G. Schiebinger, J. Shu, M. Tabaka, B. Cleary, V. Subramanian, A. Solomon, S. Liu, S. Lin, P. Berube, L. Lee et al., “Reconstruction of developmental landscapes by optimal-transport analysis of single-cell gene expression sheds light on cellular reprogramming.” BioRxiv, p. 191056, 2017.

[22] A. Forrow, J.-C. Hütter, M. Nitzan, G. Schiebinger, P. Rigollet, and J. Weed, “Statistical optimal transport via factored couplings,” arXiv preprint arXiv:1806.07348, 2018.

[23] A. Regev, S. A. Teichmann, E. S. Lander, I. Amit, C. Benoist, E. Birney, B. Bodenmiller, P. Campbell, P. Carninci, M. Clatworthy et al., “Science forum: the human cell atlas,” Elife, vol. 6, p. e27041, 2017.

[24] O. Rozenblatt-Rosen, M. J. Stubbington, A. Regev, and S. A. Teichmann, “The human cell atlas: from vision to reality,” Nature News, vol. 550, no. 7677, p. 451, 2017.

[25] “Berkeley drosophila transcription network project,” http://bdtnp.lbl.gov/Fly-Net/bioimaging.jsp.

[26] E. Levina and P. J. Bickel, “Maximum likelihood estimation of intrinsic dimension,” in Advances in neural information processing systems, 2005, pp. 777–784.

[27] J. B. Tenenbaum, V. De Silva, and J. C. Langford, “A global geometric framework for nonlinear dimensionality reduction,” science, vol. 290, no. 5500, pp. 2319–2323, 2000.

[28] “Berkeley drosophila genome project,” http://insitu.fruitfly.org/cgi-bin/ex/insitu.pl.

[29] J.-D. Benamou, G. Carlier, M. Cuturi, L. Nenna, and G. Peyré, “Iterative bregman projections for regularized transportation problems,” SIAM Journal on Scientific Computing, vol. 37, no. 2, pp. A1111–A1138, 2015.

[30] R. Sinkhorn, “Diagonal equivalence to matrices with prescribed row and column sums,” The American Mathematical Monthly, vol. 74, no. 4, pp. 402–405, 1967.

